# *Plasmodium* Protein Kinase 2 is required for ookinete to oocyst transition, and parasite transmission by the mosquito

**DOI:** 10.64898/2026.03.27.714672

**Authors:** Sarah L. Pashley, Molly Hair, Chiamaka V. Ukegbu, Mohammad Zeeshan, Akancha Mishra, Declan Brady, Sue Vaughan, Carla Pasquarello, Anthony A. Holder, Alexandre Hainard, David S. Guttery, George K. Christophides, Dina Vlachou, Pushkar Sharma, Rita Tewari

## Abstract

*Plasmodium* spp., the parasites that are the causative agents of malaria, encode a repertoire of divergent protein kinases that coordinate essential processes including cell division and host cell invasion, yet the functions of many kinases are poorly defined. *Plasmodium* Protein Kinase 2 (PK2) is essential for asexual blood-stage proliferation and has been implicated in *P. falciparum* merozoite invasion of red blood cells. However, its role in the sexual stages of the *Plasmodium* life cycle responsible for transmission is unknown. Here, using live cell imaging, functional analyses, ultrastructure microscopy and phosphoproteomics, we demonstrate that PK2 has a significant role in the *Plasmodium berghei* life cycle in the mosquito. We show that PK2 is expressed in merozoites, ookinetes and sporozoites - the invasive stages of the parasite life cycle. A conditional knockdown approach revealed that PK2 is required for the ookinete to oocyst transition in the mosquito midgut, potentially associated with altered microneme positioning’. Using haemocoel injection to bypass the midgut barrier revealed that PK2 is also required for sporozoite development after midgut invasion. Following PK2 knockdown, global proteome abundance was largely unaffected at 24 h post activation, whereas phosphoproteomics identified changes in phosphorylation of proteins linked to midgut traversal, parasite architecture, and gene regulation. These studies provide insight into the importance of PK2 function in *Plasmodium* sexual stages and parasite transmission through the mosquito, highlighting its essential function during the three invasive stages of the parasite’s life cycle.

## Introduction

Malaria is a major global public health burden, with an estimated 263 million cases and 597,000 deaths in 2023, mainly children under 5-years old (WHO, 2024). Despite the availability of multiple antimalarial therapies that target the *Plasmodium* parasite, the continued emergence and spread of drug-resistant parasites threatens control and elimination efforts [1]. Hence, the identification of novel therapeutic targets and parasite-selective drugs remains a priority. Protein kinases regulate signalling networks fundamental to parasite proliferation, development, and transmission, and therefore have attracted considerable interest as such targets [2].

The *Plasmodium* life-cycle includes asexual replication in the mammalian host and sexual development within the mosquito vector. Developmental transitions involve extensive genome replication by both mitosis and meiosis, and the generation of specialised invasive cells [3–6]. Three morphologically distinct invasive forms: merozoites, ookinetes and sporozoites are responsible for infection, replication, and transmission in mammalian host and mosquito vector stages [3, 7, 8]. Following sporozoite injection by an infected mosquito, parasites multiply in liver hepatocytes to produce merozoites, which invade erythrocytes. Cyclic propagation of merozoites and invasion of new erythrocytes during asexual blood-stage schizogony [6], results in the pathogenesis associated with the disease malaria. A subpopulation of cells differentiates into gametocytes in the bloodstream and following their ingestion by a mosquito in a blood meal, these gametocytes are activated and undergo gametogenesis. Following gamete fertilisation, the zygote differentiates (over VI stages) into an ookinete in the lumen of the mosquito midgut. The ookinete is a highly motile and invasive cell that traverses the midgut epithelium and establishes an oocyst. Over approximately 2-3 weeks, the oocyst undergoes prolonged endomitosis (termed sporogony) to produce another invasive stage: the sporozoite [9]. Sporozoites migrate to the mosquito salivary glands and are transmitted to the mammalian host in a bloodmeal. Host cell or tissue invasion by merozoites, ookinetes and sporozoites is facilitated by the polarised morphology of the cell, an actomyosin motor, and an apical secretory apparatus variously comprised of micronemes, rhoptries, and dense granules, which are subcellular organelles that secrete proteins required for motility, attachment, and host-cell entry[3].

Protein kinases (PKs) coordinate signalling pathways in parasite differentiation and host-cell invasion, and yet a substantial fraction of the *Plasmodium* kinome is poorly characterised. *Plasmodium* spp. encode approximately 60 to 90 protein kinases (depending on the species [10, 11]), many of which are highly divergent from their human orthologues, supporting their potential as tractable targets for therapeutic intervention [2, 12]. A systematic functional analysis of the *Plasmodium berghei* kinome identified 66 PKs, and of these 23 were dispensable for blood-stage proliferation in the mammalian host. Of these 23, 13 were essential for sexual stages and transmission through the mosquito. The remaining PKs were either redundant or had essential roles at other stages of parasite development in the mammalian host [10].

Here, we focus on Protein Kinase 2 (PK2), a kinase previously identified as essential for blood-stage development in *P. berghei* and *P. falciparum* [10, 13]. Reports differ regarding whether *P. falciparum* (Pf) PK2 is activated by Ca²⁺/calmodulin in vivo [14, 15], but it shares ∼55% sequence similarity with human CaMKIδ. PfPK2 is required for microneme secretion in merozoites [14], and in conditional PfPK2 knockdown parasites, the microneme protein apical membrane antigen 1 (AMA1) fails to translocate to the merozoite surface following parasite egress from the infected erythrocyte, remaining within micronemes. Consequently, merozoites attach to erythrocytes but are unable to complete invasion, and AMA1 abundance in culture supernatants is markedly reduced [14].

Although PK2 function in asexual blood stages has been defined, any role during sexual development and parasite transmission is currently unknown. Using the rodent malaria parasite *P. berghei* (Pb), here we show that PbPK2 is expressed in two additional invasive forms: ookinetes and sporozoites. Conditional depletion of PbPK2, using a gene promoter-swap strategy, ablated oocyst formation and blocked transmission. This phenotype was associated with disruption of ookinete apical microneme positioning, a process required for ookinete motility and invasion of the mosquito midgut epithelium.

## Results

### PbPK2-GFP is located at two foci within asexual blood stages

To assess the expression and subcellular location of PbPK2, we generated a PK2–GFP transgenic *P. berghei* line with GFP fused in-frame to the C-terminus of PK2 (PBANKA_1453400), expressed from the endogenous *pk2* locus (Supplementary Fig. 1A). Successful genomic integration of GFP sequence was confirmed by diagnostic PCR (Supplementary Fig. 1B), and Western blot confirmed expression of a 58kDa tagged protein of the predicted size in schizont lysates (Supplementary Fig. 1C). Using this parasite line, first we examined the expression and location of PbPK2 in asexual blood stages using live cell fluorescence imaging. PK2-GFP formed discrete puncta, often including two prominent foci, with at least one clear focus in proximity to the nucleus in all stages (ring, trophozoite and schizont) (Fig. 1A).

**Figure 1:**
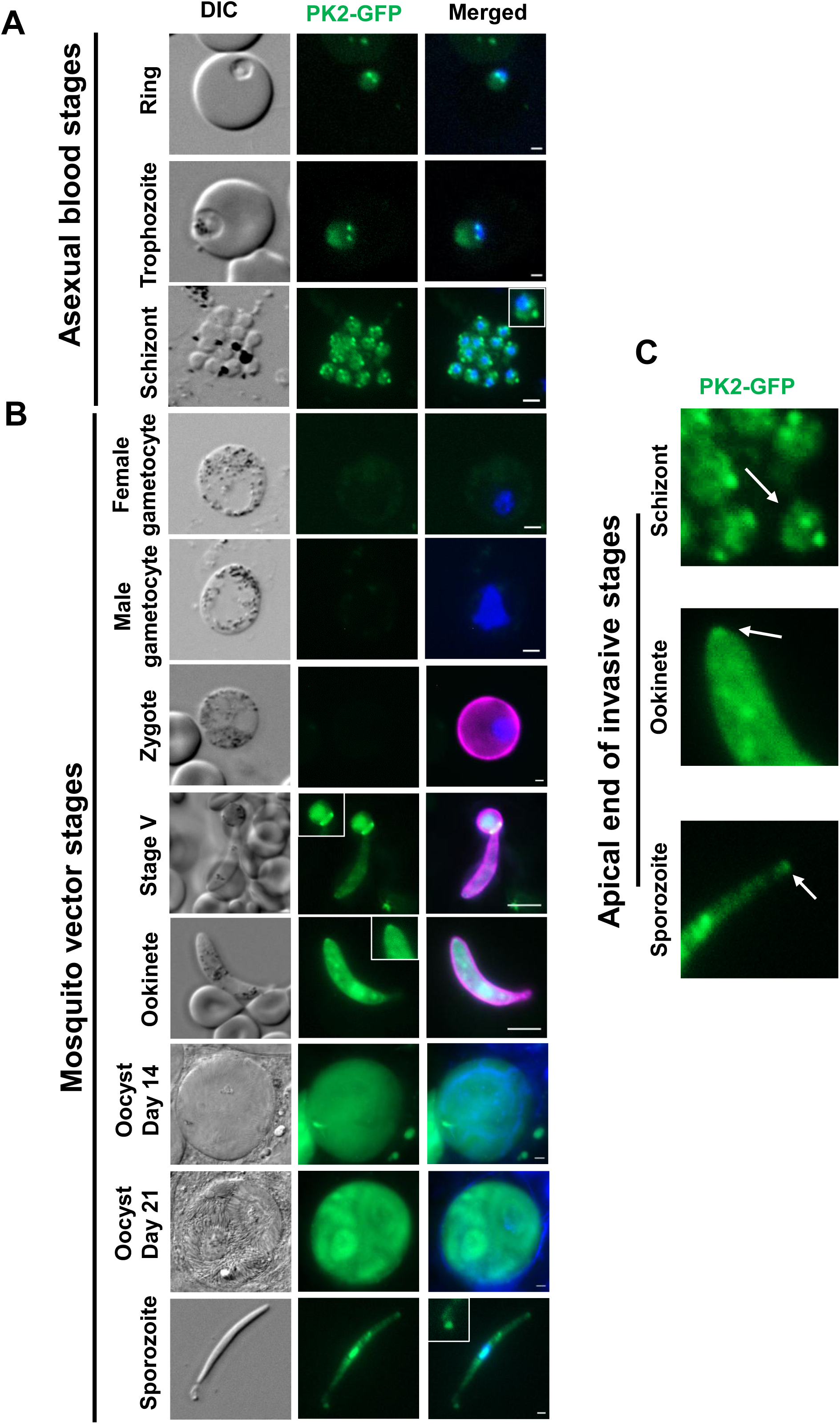
PK2-GFP is expressed in all invasive stages of the *P. berghei* life cycle. **(A)** Live cell imaging of PK2-GFP in blood stage parasites: ring, trophozoite and schizont stages. Panels are Differential Interference Contrast (DIC), GFP fluorescence (PK2-GFP) and PK2-GFP merged with Hoechst staining for nuclei. White box in schizont image contains a zoom of an individual merozoite. Scale bars, 2µm. **(B)** Live cell imaging of PK2-GFP in female and male gametocyte, zygote, stage V and mature ookinete, oocyst at day 14 and oocyst at day 21, and sporozoite. Panels are Differential Interference Contrast (DIC), GFP fluorescence (PK2-GFP), and merge of PK2-GFP, Hoechst and the 13.1 antibody (magenta) for zygote and ookinete stages. White boxes indicate zoomed regions within the corresponding image. Scale bars, 2 µm gametocytes and zygote; 5 µm stage V and mature ookinete, oocysts. White boxes indicate zoom regions within image panels. **(C)** PK2-GFP fluorescence imaging of the apical end of three invasive stages: a merozoite within a schizont, an ookinete and a sporozoite. The white arrows indicate the apical tip of the cell.

### PbPK2-GFP is expressed at various stages within the mosquito vector, with a single focus at the apical end of invasive stages

We examined PK2-GFP in mosquito stage parasites by live cell imaging to look for expression in the sexual stages of the life cycle, particularly the invasive ookinetes and sporozoites. No detectable GFP signal was observed in either male or female gametocytes (Fig. 1B), or in zygotes and during early ookinete differentiation (stages I–III) (Fig. 1B). In contrast, PK2–GFP was visible in stage IV ookinetes, increasing markedly by stage V (Fig. 1B). At stage V, the signal resolved into two prominent foci connected by a linear structure, resembling the position and morphology of the developing contractile ring. In mature ookinetes (stage VI), PK2–GFP remained readily detectable, but in a more distributed pattern along the parasite body, with stronger foci frequently enriched toward the apical end. About 65% of imaged ookinetes had PK2-GFP concentrated at the apical tip (Fig. 1B).

To define the location of PK2-GFP more clearly at the apical tip of ookinetes, we performed a genetic cross of PK2-GFP with a Myosin A–mCherry reporter line [16]. PK2–GFP and MyoA–mCherry blood stage parasites were grown in separate mice, and then combined in ookinete medium to enable gametogenesis, fertilisation and recombination. After 24 hours, live cell imaging was used to examine these cultures, and several ookinetes expressing both PK2-GFP and MyoA-mCherry were identified, indicative of successful recombination. There was a concentration of PK2-GFP at the apical end of the ookinete, subjacent to MyoA-mCherry that lined the periphery of the ookinete but with an increased intensity at the apical tip (Supplementary Fig 1D). These observations were validated at higher spatial resolution using three-dimensional structured illumination microscopy (3D-SIM) of purified, fixed ookinetes.

3D-SIM confirmed the enrichment of PK2–GFP at the apical tip (Supplementary Fig 1E), with PK2–GFP subjacent to the MyoA-mCherry-label (Supplementary Fig 1F).

To investigate PK2-GFP expression during sporogony, we allowed *Anopheles stephensi* mosquitoes to feed on mice infected with PK2-GFP parasites and analysed oocyst development on day 7, 14 and 21 post-blood meal. PK2–GFP was not detectable in day 7 oocysts (data not shown) or in day 14 oocysts prior to sporulation (Fig. 1B). In contrast, PK2–GFP was evident in day 14 oocysts that had initiated sporulation and was strongly expressed in fully sporulated oocysts at day 21 (Fig. 1B). We also isolated sporozoites from homogenised midguts and salivary glands for imaging at day 21 post-feeding. PK2–GFP was detectable in both midgut and salivary gland sporozoites (Fig. 1B), with a dispersed location but with consistent enrichment at the apical tip of the cell, mirroring the pattern observed in ookinetes. Collectively, these GFP fluorescence data indicate that PK2-GFP is predominantly located at the apical end of all the invasive stages of the parasite life cycle (Fig. 1C).

### Expansion microscopy of invasive stage parasites reveals an accumulation of PK2-GFP at the apical pole of the cell

To locate PK2 at higher spatial resolution, we analysed PK2–GFP schizonts by 3D structured illumination microscopy (3D-SIM). Consistent with the live cell imaging, PK2–GFP was predominantly located at two foci outside the nucleus (Fig. 2A). We then applied ultrastructure expansion microscopy (U-ExM) with staining of tubulin and GFP to assess the proximity of PK2-GFP to microtubules. The PK2–GFP signal showed limited overlap with that of the intranuclear spindle microtubules, indicating that PK2 is extra-nuclear and not a major component of the mitotic spindle apparatus in schizonts (Fig. 2B).

**Figure 2:**
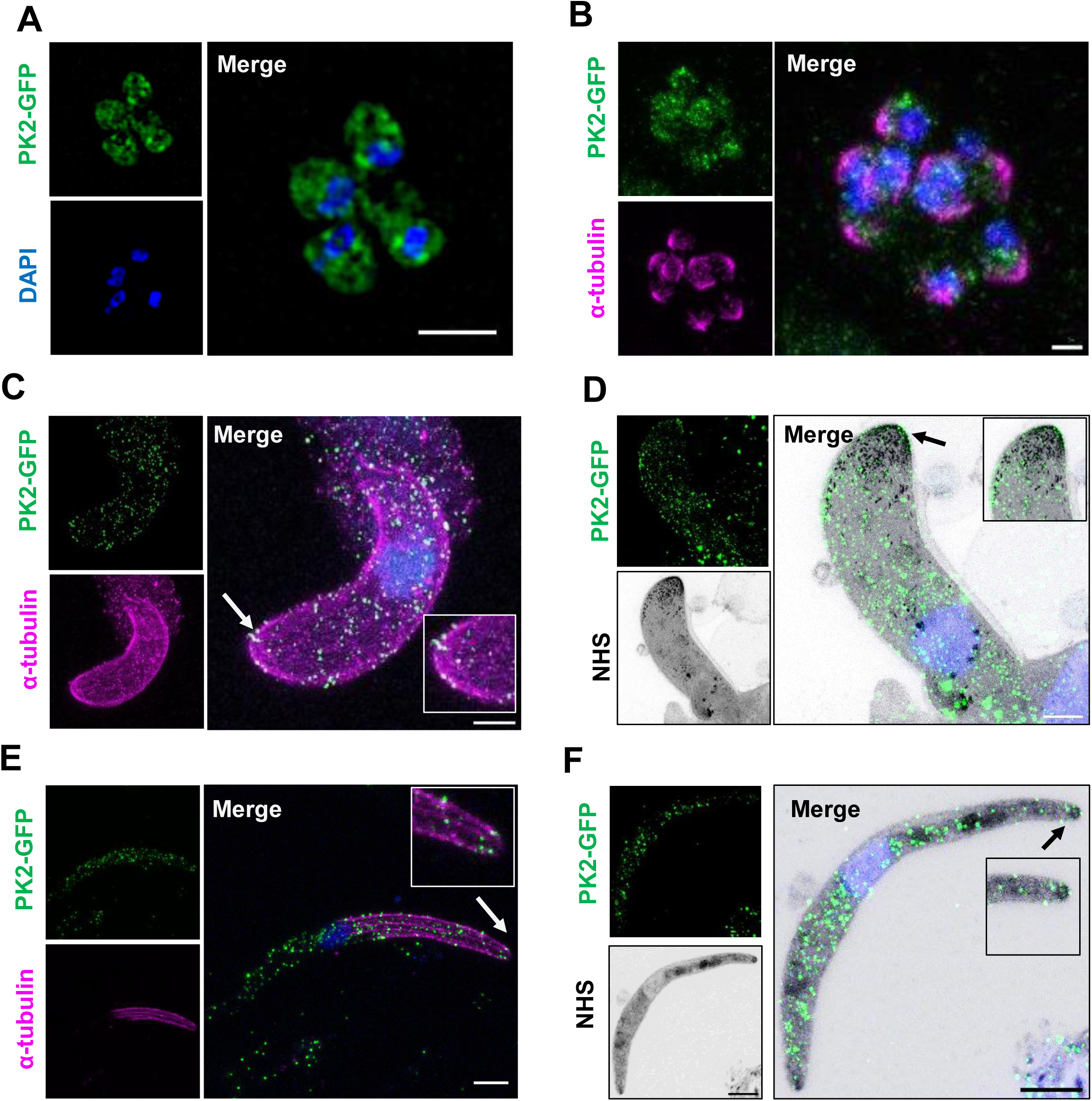
The location of PK2 in invasive parasite stages revealed using expansion microscopy. **(A)** 3D-SIM imaging of fixed PK2-GFP schizonts. Panels are PK2-GFP fluorescence (green), DAPI DNA stain (blue) and merged. Scale bar, 5 µm. **(B)** U-ExM imaging of fixed PK2-GFP schizonts labelled with GFP (green) and α-tubulin (magenta). In the merge panel the two signals are merged together with Hoechst DNA staining (blue). Scale bar, 2 µm. **(C)** U-ExM imaging of a fixed PK2-GFP ookinete labelled with GFP (green) and with α-tubulin (magenta). In the merge panel the two signals are merged, together with Hoechst DNA staining (blue). The apical end of the cell is indicated with the white arrow and the white box indicates the zoomed area within the image. Scale bar, 2 µm. **(D)** U-ExM imaging of fixed PK2-GFP ookinetes labelled with GFP (green) and with NHS-ester (grey). In the merge panel the two signals are merged, together with Hoechst DNA staining (blue). The black arrow indicates the apical end of the cell and the black box indicates the zoomed area within the image. Scale bar, 2 µm. **(E)** U-ExM imaging of a PK2-GFP sporozoite labelled with GFP (green) and with α-tubulin (magenta). In the merge panel the two signals are merged, together with Hoechst DNA staining (blue). The white arrow indicates the apical end of the cell and the white box indicates the zoomed region within the image panel. Scale bar, 2 µm. **(F)** U-ExM imaging of a PK2-GFP sporozoite labelled with GFP (green) and with NHS-ester (grey). In the merge panel the two signals are merged, together with Hoechst DNA staining (blue). The black arrow indicates the apical end of the cell, and the black box indicates the zoomed area within the image. Scale bar, 2 µm.

Next, we examined PK2–GFP distribution in free merozoites using U-ExM with GFP detection alongside NHS-ester labelling of prominent subcellular apical structures such as rhoptries that are involved in invasion. Multiple points of PK2–GFP signal were observed, typically including two larger and more intense foci (Supplementary Fig. 1E). Although the relative positions of these foci varied, a subset of merozoites had a prominent focus close to the rhoptry neck region (Supplementary Fig. 1E). Together, these observations suggest that PK2-GFP is largely concentrated in distinct foci within blood stage parasites, with a fraction frequently enriched toward the apical end of the cell.

To investigate further the location of PK2-GFP in mature ookinetes, we used U-ExM with anti-tubulin and anti-GFP staining on fixed samples. GFP staining was punctate and broadly distributed throughout the ookinete, similar to the pattern in live-cell imaging (Fig. 2C). However, in a subset of parasites we observed clusters of PK2–GFP foci at the apical tip of the cell (Fig. 2C). Using U-ExM with anti-GFP and NHS-ester, ookinetes were examined to determine whether these foci co-localised with subcellular structures involved in invasion. PK2–GFP foci were observed at the apical end, but they did not consistently co-localise with NHS-ester–positive structures that resembled micronemes (Fig. 2D).

Finally, to confirm PK2-GFP expression in sporozoites, we used U-ExM on fixed sporozoites stained with anti-tubulin and anti-GFP antibodies. As observed in live imaging, PK2–GFP was largely dispersed, but some additional foci were present at the apical tip in a subset of cells (Fig. 2E). PK2–GFP was not confined to and did not strongly co-localise with NHS-ester labelled microneme- or rhoptry-associated compartments at the sporozoite tip (8 cell imaged) (Fig. 2F).

### Conditional knockdown of PbPK2 revealed a role for PK2 in the ookinete to oocyst transition and parasite transmission

Our observation that PK2-GFP is expressed in invasive ookinete and sporozoite stages in the mosquito led us to examine the function of PK2 in these stages. We created a conditional knockdown *P. berghei* mutant (henceforth termed *pk2ptd*) that placed the *pk2* gene under the control of the *ama1* promoter (PBANKA_0915000), which has low transcript levels in *P. berghei* gametocytes [17] (Supplementary Fig. 2A). Correct integration of the *ama1* promoter was confirmed by diagnostic PCR (Supplementary Fig. 2B). qRT-PCR confirmed that *pk2* transcript levels were significantly reduced by 70% in activated *pk2ptd* gametocytes and by 93% in *pk2ptd* ookinetes, respectively (Supplementary Fig. 2C & D).

Male gamete production (assessed by the formation of exflagellation centres) was unaffected in the *pk2ptd* line compared to WT-GFP controls (Fig. 3A), and fertilisation and zygote to ookinete conversion were also unaffected (Fig. 3B). However, *pk2ptd* oocyst development in mosquitoes was almost completely ablated. No *pk2ptd* oocysts were observed at day 7 post infection, only 11 were detected at day 14 (Fig. 3C), and only 1 was detected at day 21, respectively, while hundreds of oocysts were present at each time point in comparable mosquitoes fed WT-GFP parasites. The few *pk2ptd* oocysts were significantly smaller than WT-GFP oocysts (Fig. 3D). To confirm the absence of any viable oocyst or sporozoite, infected mosquitoes were allowed to feed on naïve mice. While mosquitoes were able to transmit the WT GFP parasite, and blood stage infection was observed on average 5 days later (Fig. 3E), mosquitoes infected with *pk2ptd* parasites failed to infect susceptible mice,

**Figure 3:**
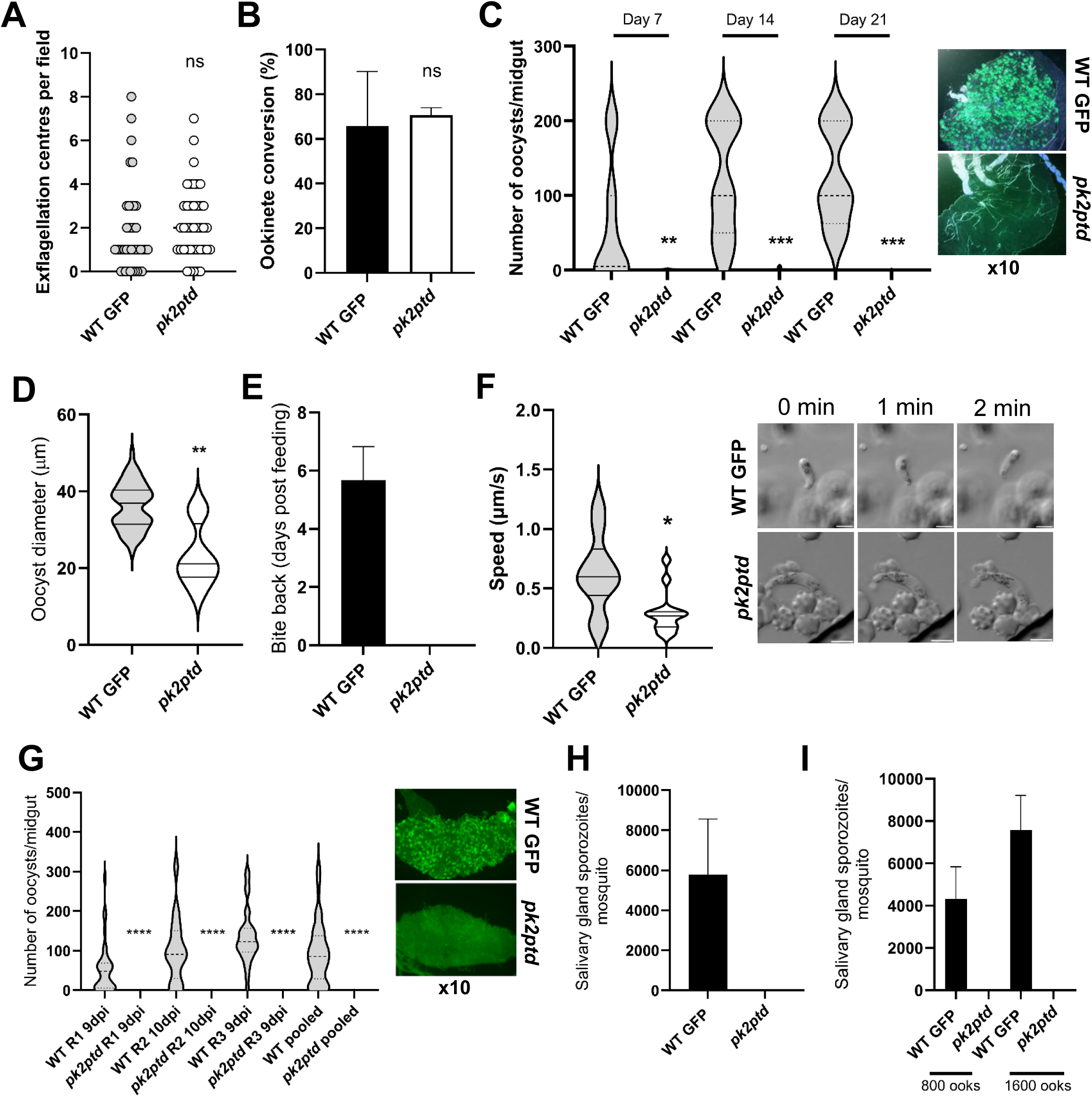
PbPK2 is required for the ookinete to oocyst transition and sporozoite development. **(A)** Exflagellation centres per field at 15-min post activation. n = 3 independent experiments (>20 fields per experiment).. **(B)** Percentage ookinete conversion. *n* = 3 independent experiments (>100 cells). Error bar, ± SEM **(C)** Total number of GFP-positive oocysts per infected mosquito in *pk2ptd* compared to WT GFP parasites at 7-, 14- and 21-days post infection. Mean ± SEM. n = 3 independent experiments. **P <0.01; ***P <0.001. Representative images of WT GFP and *pk2ptd* infected midguts at 10x magnification. **(D)** Diameter of individual *pk2ptd* and WT GFP oocysts (μm) at 14-days post-infection. Horizontal line indicates the mean from three independent experiments (20 mosquitoes for each of *pk2ptd* and WT GFP). **P < 0.01. **(E)** Bite-back experiments for transmission of WT GFP and *pk2ptd* parasites from mosquito to mouse. Infected mosquitoes were fed on mice and then the mice were monitored for day of patent blood stage parasitaemia indicative of successful infection. N = two independent experiments. **(F)** Velocity of individual WT GFP or *pk2ptd* ookinetes from 24 hr cultures measured over 2 min, and shown in the violin plot. Bar = arithmetic mean. *P<0.05. Representative frames from time-lapse movies of WT GFP (upper panels) and *pk2ptd* (lower panels) ookinetes in Matrigel at 0, 1 and 2 minutes. Scale bars = 5 μm. **(G)** Number of GFP-positive oocysts per mosquito infected with *pk2ptd* and WT GFP parasites at 9 or 10-days post infection. Mean ± SEM. n = 3 independent experiments. ****P <0.0001. Representative images of midguts infected with WT GFP and *pk2ptd* parasites at 10x magnification are shown. **(H)** Number of sporozoites per mosquito in 21-day postinfection salivary glands for WT GFP and *pk2ptd* parasite lines. Three independent experiments, n = 20 mosquitoes for each replicate. **(I)** Salivary gland sporozoites per mosquito following direct injection of WT GFP or *pk2ptd* ookinetes into the mosquito hemocoel (inocula of 800 or 1600 ookinetes). Mean ±SEM. N = 2 independent experiments per inoculum.

We analysed ookinete motility (Patzewitz *et al*., 2013) and found that gliding motility in *pk2ptd* ookinetes was significantly reduced compared to that of WT parasites (Fig. 3F; Supp. Videos 1 and 2), indicating that their motor function is impaired and suggesting that this defect may be the cause of the ablation of oocyst development in *pk2ptd* lines, potentially by preventing invasion of the midgut epithelium. In an independent insectary (Imperial College), mosquitoes infected with the same lines showed ablation of *pk2ptd* oocyst development, with no oocysts detected 9 or 10 days after infection (Fig. 3G), and no sporozoites recovered from mosquito salivary glands 21 days post infection (Fig. 3H). In contrast, with the WT GFP control line hundreds of oocysts were detected in mosquitoes (Fig. 3G), resulting in several thousand sporozoites per mosquito (Fig. 3H).

To determine whether the lack of *pk2ptd* oocysts and sporozoites was just due to a defect in invasion of the midgut epithelium, we bypassed the midgut barrier by injecting either 800 or 1600 ookinetes directly into the haemocoel of *A. stephensi* mosquitoes and analysed salivary glands for sporozoites 21 days post-injection [18]. No sporozoites were observed in mosquitoes infected with *pk2ptd* ookinetes, whereas the WT GFP control line produced several thousand sporozoites per mosquito (Fig. 3I), suggesting that while *pk2ptd* ookinete motility is significantly affected and PK2 may have a significant role in midgut invasion by ookinetes, it has an additional role in sporozoite development independent of ookinete motility.

### Ultrastructural studies of *pk2ptd* ookinetes showed that PK2 is important for microneme formation and distribution

We used U-ExM and transmission electron microscopy (TEM) to look for ultrastructural defects in *pk2ptd* ookinetes that may be responsible for the impaired invasion phenotype. U-ExM with anti-tubulin antibody and NHS-ester labelling of protein-dense compartments showed *pk2ptd* mature ookinete morphology to be similar to that of the WT GFP control (Fig. 4A). In contrast, a quantitative analysis of apical micronemes/granules revealed a significant reduction in *pk2ptd* parasites (mean 32) relative to WT GFP (mean 76, *P* <0.0001), with microneme-like structures frequently present away from the apex and toward the basal region in *pk2ptd* parasites (Fig. 4A). This spatial shift is consistent with a defect in the apical targeting or retention of micronemes during ookinete maturation.

**Figure 4:**
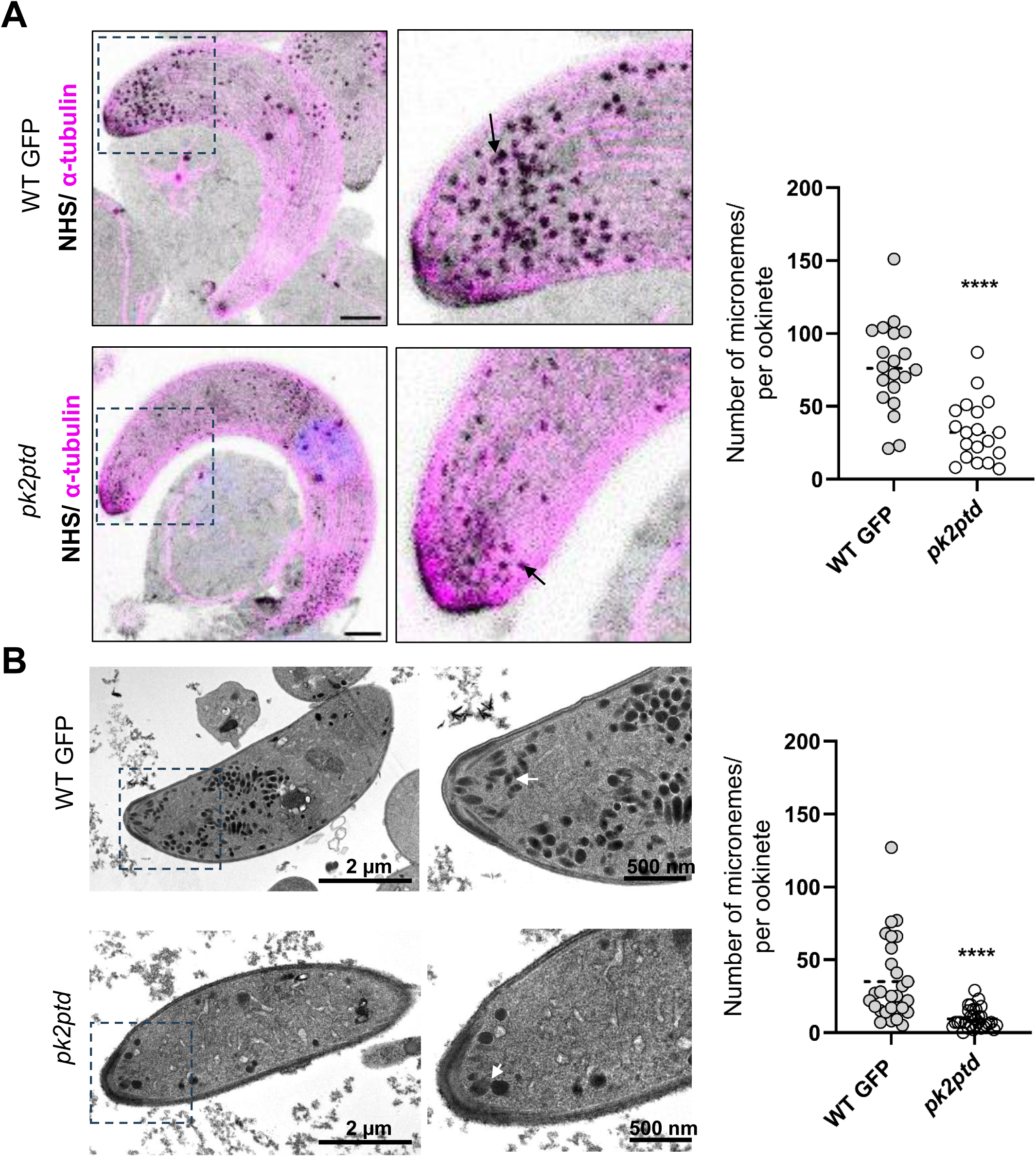
*pk2ptd* ookinetes have altered distribution of apical micronemes. **(A)** U-ExM imaging of WT GFP and *pk2ptd* ookinetes stained with α-tubulin (magenta), NHS-ester (grey) and Hoechst (blue). Scale bars, 2 µm. Black dashed boxes on the lefthand panels indicated the region of the images shown at higher magnification in the right-hand panels. Right panel: Number of micronemes at the apical end in WT GFP vs *pk2ptd* ookinetes identified by NHS-ester staining and quantified using ImageJ. **(B)** TEM imaging of WT GFP and *pk2ptd* ookinetes. The black dashed boxes on the left-hand panels indicate the region of the images shown at higher magnification in the right-hand panels. Scale bar size indicated in images. Right panel: Number of micronemes at the apical end in WT GFP vs *pk2ptd* ookinetes from TEM images, quantified using ImageJ.

To support these U-ExM observations, we used TEM with both WT GFP and *pk2ptd* mature ookinetes. TEM confirmed that the overall cellular organisation of *pk2ptd* ookinetes was similar to that of WT GFP parasites (Fig. 4B), and that there was a significant depletion of apical micronemes (*pk2ptd* mean 9.5 versus WT GFP mean 35.1, *P <*0.0001), with an increased number of micronemes toward the basal end of *pk2ptd* parasites (Fig. 4B).

Because microneme abundance and positioning were altered, we asked whether the transcript levels of key microneme-associated invasion proteins were significantly altered in *pk2ptd* ookinetes. qRT–PCR analysis of 24 h ookinete cultures revealed a modest, but non-significant increase in CTRP transcription, whereas SOAP, CHT1, P28, and CelTOS transcript levels were similar in WT GFP and *pk2ptd* parasites (Supplementary Fig. 2E, F). These data suggest that the microneme mislocalisation phenotype of *pk2ptd* ookinetes is unlikely to be driven by a widespread transcriptional downregulation of genes for canonical microneme cargo.

### Differential phosphorylation of multiple proteins involved in midgut invasion and gene expression in *pk2ptd* parasites

To examine how PK2 reduction impacts total protein abundance and phosphorylation in mature ookinetes, we performed quantitative phosphoproteomic analysis (with phosphopeptide enrichment) comparing WT GFP and *pk2ptd* parasites at 24 hours post-activation (hpa) (Fig. 5). Using thresholds of log_2_ FC ≥0.58 or ≤-0.58 and *P* ≤ 0.05, the proteomic analysis revealed few changes in overall protein abundance, with only three proteins significantly reduced and none significantly increased in *pk2ptd* parasites (Fig. 5A). SOAP, CHT1 and PSOP20 were decreased in the *pk2ptd* line. These data indicate that the ookinete proteome at 24 hpa is largely preserved, despite the depletion of PK2. In contrast, the phosphoproteomic analysis identified significant changes in 51 phosphopeptides in *pk2ptd* parasites (15 decreased; 36 increased; Fig. 5B; Supplementary Table 1, 2). These sites of differential phosphorylation mapped to proteins important in inner membrane complex (IMC)/cytoskeletal organisation, host cell interactions/secreted factors, and nuclear regulation (Fig. 5C). Notably, several IMC/cytoskeletal-associated proteins had reduced phosphorylation in *pk2ptd* parasites, including ALV7 and the zinc finger protein Pb103, whereas several phosphosites on IMC1i were hyperphosphorylated. Phosphorylation of host interacting proteins such as EMAP1 [19] was reduced, whereas the perforin-like protein PLP4, which has previously been shown to be necessary for ookinete traversal of the midgut epithelium [20, 21], was found to be hyperphosphorylated in *pk2ptd* ookinetes. Changes in the phosphorylation of proteins linked to chromatin/DNA repair and transcriptional regulation included NOT1-G. Together, these data indicate that *pk2ptd* parasites with depleted PK2 have significantly altered phosphorylation of proteins important in ookinete architecture, host interaction and midgut traversal, despite only modest alterations in overall protein abundance.

**Figure 5:**
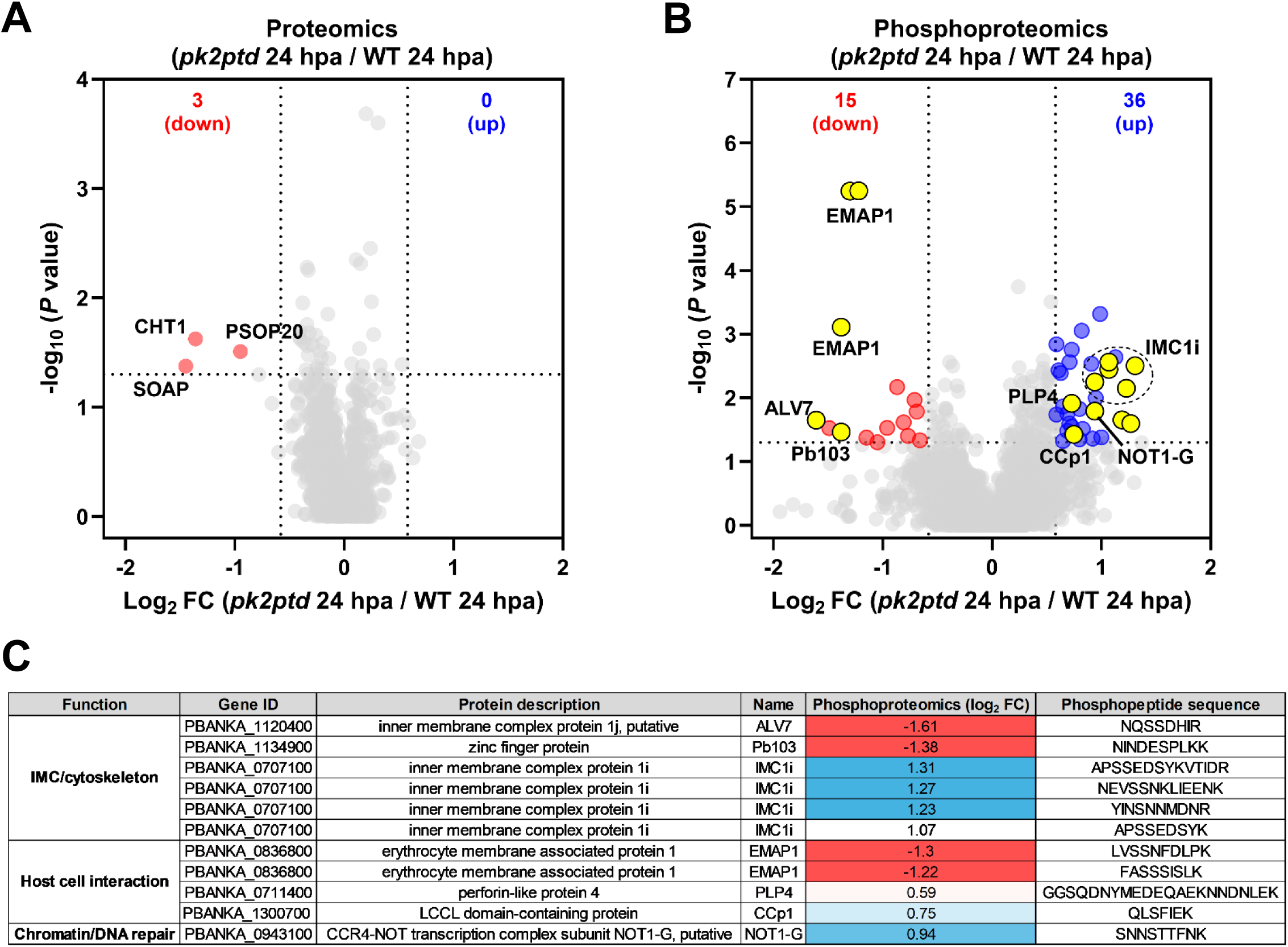
PK2 depletion in ookinetes at 24 hpa has little effect on total protein abundance but drives changes in phosphorylation. **(A)** Volcano plot comparison of total proteome from *pk2ptd* and WT GFP parasites at 24 hours post activation (hpa). Points are quantified proteins plotted as log_2_ fold-change (*pk2ptd*/WT) versus –log_10_ (*P* value); significantly decreased proteins are highlighted (orange) and labelled (SOAP, CHT1, PSOP20). **(B)** Volcano plot comparison of phosphoproteome changes at 24 hpa ookinetes from *pk2ptd* and WT GFP lines. Phosphosites significantly decreased (red) or increased (blue) in abundance are indicated, with examples labelled (e.g., EMAP, CCp1, PLP4, IMC1i, NOT1-G) in yellow. **(C)** Summary table of selected differentially phosphorylated proteins grouped by function, showing gene ID, protein annotation, log_2_ fold-change, and corresponding phosphopeptide sequence. Vertical and horizontal dashed lines in (A and B) indicate fold-change and significance thresholds used for hit calling.

## Discussion

Protein kinases are successfully targeted in the treatment of many diseases such as cancer [22] and efforts are being made to develop therapeutics that target *Plasmodium* kinases [12]. PK2 function has been investigated in asexual blood stage parasites, but little was known about its importance in transmission, so we investigated its role in the sexual stages of the life cycle. A previous study had begun to characterise *P. falciparum* PK2, showing that it was required for merozoite invasion [14]. PK2 was found in distinct foci in schizonts; however, these did not colocalise well with components of the invasive machinery, such as micronemes or rhoptries. We have also found PK2 at discrete foci in *Plasmodium berghei* blood stages, not only in schizonts but also in rings and trophozoites.

Live cell imaging of PK2-GFP parasites revealed that there is no detectable expression in male or female gametocytes. However, post-fertilisation, we found PK2 expression begins around ookinete development stage IV, with an interesting pattern reminiscent of the contractile ring, similar to that of Myosin-F or Kinesin-20 [16, 23]. PK2 was present in mature ookinetes, diffusely distributed throughout the cytoplasm with some foci and a concentration at the apical tip. PK2 was also expressed in sporozoites and again with a particular concentration at the apical tip, similar to the location of Myosin B and other conoid associated proteins[24–27]. Together, these experiments showed that PK2 is expressed in all the invasive stages of the *P. berghei* life cycle, suggesting that it is involved as a common component of the invasion process in all three stages, not just in merozoites as described previously [14].

Of particular interest is that PK2 is required for the ookinete to oocyst transition in the mosquito midgut. The conditional knockdown approach that reduced PK2 transcript levels in the *pk2ptd* ookinete, produced a mutant parasite unable to establish an infection in the mosquito midgut, despite numbers of motile ookinetes similar to those of GFP WT parasites. This phenotype is like that of a CBP-O knockout mutant and, interestingly, CBP-O is also located at the ookinete apical tip [24]. Strikingly, even after direct injection of *pk2ptd* ookinetes into the mosquito haemocoel, the parasites were still unable to form oocysts and produce sporozoites. This indicates that not only is PK2 required for ookinete invasion of the midgut epithelium, but it also has an additional later role in sporozoite development. This phenotype is quite different to that of SHLP1 mutants, which have a similar mislocalization of micronemes but with a defect that was solely limited to ookinetes [18]. A similar phenotype was observed previously in a PIMMS43 mutant: direct injection of these ookinetes into the haemocoel also failed to produce sporozoites [28]. Although, our PK2-GFP imaging data show that PK2 is expressed in sporulated oocysts, it is possible that PK2 has a role in the later stages of sporozoite development, after DNA replication in the formation of the sporoblast. Clearly, PK2 is essential since oocysts are not viable without it. Future experiments are needed to characterise the role of PK2 during sporozoite development and whether it is also important for sporozoite invasion. PK2 is essential for the parasite transmission stages and has a critical function in ookinete to oocyst transition and sporozoite development.

The phosphoproteomic analysis provides a mechanistic insight into how PK2 may have key roles in transmission-stage development. Several proteins known to be essential for oocyst development or sporogony were differentially phosphorylated in *pk2ptd* ookinetes. In particular, the LCCL-domain containing protein CCp1, whose disruption ablates oocyst development [29], was differentially phosphorylated. The phosphorylation/dephosphorylation (hyper- versus hypophosphorylation) suggests that PK2 acts both through direct substrate phosphorylation and through activation of phosphatase activity. The perforin-like protein PLP4, implicated in midgut traversal [20, 21] and IMC1i, implicated in ookinete development [30], were hyperphosphorylated; whereas kinesin-8X, which is required for oocyst development and sporogony [31], was hypophosphorylated. Collectively, these changes support a model in which PK2 regulates a phosphorylation/dephosphorylation network spanning invasion-associated functions and developmental programmes required for ookinete to oocyst transition and parasite transmission. This profile differs from that resulting from the perturbation of PfPK2 in asexual blood stages, which was linked to cGMP/Ca²⁺-associated invasion signalling and changes in phosphorylation in invasion-related pathways [14], highlighting the context-specific functions of PK2-regulated networks in different life-cycle invasive stages.

Our observation that *pk2ptd* mutant ookinetes had an altered location of micronemes is similar to that observed previously in a SHLP1 knockout mutant [18]. The release of microneme contents is necessary for ookinete invasion of the mosquito midgut[18, 32], and the secreted proteins have several functions in ookinete adhesion, motility and membrane degradation. For example, CTRP is required for ookinete motility and its gene deletion results in a failure to establish midgut infection [33]. CHT1 is required to degrade the peritrophic matrix of the gut and its gene disruption significantly reduces midgut infection [34, 35]. In contrast to other invasive stages of the life cycle, ookinetes lack both rhoptries and dense granules and invasion relies mainly on micronemes [36, 37].

Normally, micronemes bud off from the ER-Golgi secretory pathway and are transported along microtubules towards the apical end of the cell, eventually docking with rhoptry tips at the time of release [38–40]. Mis-localisation of micronemes could indicate transport defects and may also indicate that they are either unable to secrete components required for invasion or are able to secrete but the components are in the wrong sub-cellular compartment, potentially explaining why so few oocysts are formed [18, 36]. The latter argument may explain why the ookinetes are motile but unable to invade, with other components such as CTRP being essential for ookinete motility [33]. It is also possible that different types of microneme may exist [41]. It will be important to determine whether the different types of microneme exist in ookinete and sporozoites and whether they have functional differences. Our observations on the importance of micronemes fit with findings in *P. falciparum* merozoites, showing that secretion of microneme proteins is reduced in a conditional knockout mutant of PK2 [41]. Taken together, these findings suggest that PK2 has an important function with respect to micronemes, either in ensuring their correct positioning or directly or indirectly facilitating release of their contents in all three invasive stages of malaria parasite.

Overall, our data indicate that PK2 is a protein kinase that is essential for parasite transmission through mosquitoes and has a conserved apical location and association with secretory organelle function across the different invasive forms. We propose that PK2 regulates a programme of phosphorylation/dephosphorylation that is required for correct microneme distribution and successful ookinete to oocyst transition. It has additional roles during oocyst maturation and/or initiation of sporogony. These findings support the idea of PK2-regulated pathways as attractive candidates for transmission-blocking intervention strategies.

## Materials and methods

### Ethics statement

Animal procedures were carried out in accordance with the Animal (Scientific Procedures) Act 1986 under the UK Home Office Licenses PDD2D5182; PP3589958 and PPL/PP3947717. Female outbred CD1 mice aged 6-8 weeks were used for all experiments. Mice were kept in a 12-hour light, 12-hour dark cycle at a temperature between 20 – 24°C with humidity between 40 – 60%.

### Generation of transgenic parasites

PK2-GFP was generated by amplifying a region of the *Pbpk2* gene downstream of the ATG start codon and ligating it to the p277 vector as described previously [32]. The p277 vector contains the human *dhfr* cassette, conveying resistance to pyrimethamine. Schematic representations of the endogenous gene locus, the constructs and the recombined gene locus can be found in Supplementary Fig 1A. Diagnostic PCR was performed with primers IntT302 and ol492. For all primers used in the study, see Supp. Table 3.

The *pk2ptd* parasite line was generated using a conditional knockdown strategy which places the *pk2* gene under the control of the *ama1* promoter. The knockdown construct was generated by double homologous recombination into the *Pama1* (pSS368) vector. A schematic representation of the endogenous *pk2* locus, the constructs and the recombined *pk2* locus can be found in Supplementary Fig 2A. Diagnostic PCR was performed for *pk2* gene knockdown parasites using primers IntPTD61_5 and 5′-IntPTD to determine successful integration of the targeting construct at the 5′ end of the gene locus. Primers IntPTD61_3 and 3′-IntAMA1 were used to determine successful integration for the 3′ end of the gene locus. Both constructs were transfected by electroporation [42].

### Generation of dual-tagged parasites

Blood from a mouse injected with PK2-GFP was mixed with blood from a mouse injected with MyosinA-mCherry [16]. For ookinetes, the blood from each mouse was mixed in equal volumes with ookinete media. Recombination took place in vitro and some crossed ookinetes were produced. To create the line, the mixed blood was injected into an uninfected mouse. Once parasitaemia and gametocytaemia were sufficient, *A. stephensi* mosquitoes were allowed to feed. After 21 days, a bite back experiment was performed to generate a new crossed parasite line. Blood was collected after 5 days when parasitaemia was high enough and the blood examined to find crosses.

### Purification of schizonts and gametocytes

Blood from infected mice (four days post-infection) was placed in schizont medium (RPMI-1640 medium containing penicillin-streptomycin, and heat inactivated FBS) for 24 hours at 37°C with rotation at 100 rpm. Schizonts were then purified on a 60% (in PBS) v/v nycodenz column. NycoDenz stock solution: 27.6% w/v NycoDenz in 5 mM Tris-HCl (pH 7.20), 3 mM KCl, 0.3 mM EDTA). Schizonts were then harvested from the interface and used for downstream assays.

The purification of gametocytes was achieved by injecting parasites into phenylhydrazine-treated mice [43] and enriched by sulfadiazine treatment after 2 days of infection. The blood was collected on day 4 after infection and gametocyte-infected cells were purified on a 48% v/v NycoDenz (in PBS) gradient (NycoDenz stock solution: 27.6% w/v NycoDenz in 5 mM Tris-HCl (pH 7.20), 3 mM KCl, 0.3 mM EDTA). The gametocytes were harvested from the interface and activated for 15 mins to induce exflagellation.

### Ookinete culture and purification

Blood from infected mice was cultured in ookinete medium (RPMI-1640, 20% FBS, 100 µM xanthurenic acid) for up to 24 hours to produce mature ookinetes. Ookinetes were purified using one of two methods. Either by Nycodenz column (62.5% in PBS) when the aim of the experiment was RNA extraction or by magnetic beads coated in an antibody against surface protein P28, for U-ExM or TEM experiments.

### Live cell imaging

To determine PK2 expression throughout the *Plasmodium* life cycle, small amounts of blood containing PK2-GFP parasites was mixed with PBS-DAPI and imaged to visualise blood stage parasites. Blood was also placed into schizont culture and imaged at regular intervals over a period of 24 hours. Blood was also added to ookinete media (RPMI media containing 100 μM xanthurenic acid, 1% w/v sodium bicarbonate, and 20 % v/v heat inactivated FBS) and ookinete development captured by live cell imaging. 13.1-Cy3 antibody was used to identify the P28 surface protein expressed by zygotes and ookinete stages. Oocysts and sporozoites were imaged by removing the mosquito midguts/salivary glands from the mosquito. Salivary glands were homogenised in PBS-DAPI to visualise free sporozoites. A 63x oil immersion objective was used to capture all images on a Zeiss AxioImager M2 microscope fitted with an AxioCam ICc1 digital camera.

### Parasite phenotype analysis

For analysis of male gametocyte exflagellation, mice were injected with blood containing approximately 1 x 10^6^ parasites. 4-5 days later, blood was collected and exflagellation centres were counted by activating a small volume of blood in ookinete media for 15 minutes. For ookinetes, parasite infected blood was incubated in ookinete media at 20°C for 24 hours and ookinete conversion was quantified using 13.1 antibody. To assess oocyst formation and development, 30-50 *Anopheles.stephensi* SD 500 mosquitoes were allowed to feed for 20 minutes on anaesthetised, infected mice that had comparable parasitaemia and gametocytaemia. Around 10 midguts were removed from infected mosquitoes on days 7, 14 and 21 post-feeding and imaged to assess oocyst number. Mosquito bite back experiments were performed by allowing infected mosquitoes to feed on anaesthetised, uninfected mice. Blood smears were made from these mice and days counted until the mice became positive for parasites.

### Ookinete injection experiments

3–5-day old female *Anopheles stephensi* mosquitoes were infected by microinjection of 69 nl of *in vitro* cultured ookinetes into the thorax using glass capillary needles and the Nanoject II microinjector (Drummond Scientific). Briefly, WT GFP and *pk2ptd* ookinetes were cultured as described. Blood from infected mice was collected via cardiac puncture and placed in 30 volumes of the ookinete medium (RPMI 1640 supplemented with 25 mM HEPES, 0.4 mM hypoxanthine, 0.1 mM xanthurenic acid, 24 mM of NaHCO_3_ and 100 U/ml of penicillin/streptomycin solution (Thermo Fisher Scientific)) before incubation for 24 hr at 21°C. Ookinete numbers from 22–24-hour *in vitro* cultures were counted using a standard haemocytometer and ookinete density adjusted to 1.2 or 2.4 x 10^4 per ul for a final concentration of ∼800 or 1,600 ookinetes per mosquito respectively. At 20-21 dpbi, 25 salivary glands were dissected, homogenised in PBS and salivary gland sporozoites counted using a standard haemocytometer.

### Western blotting

Purified schizonts were lysed in lysis buffer (10 mM Tris-HCl [pH 7.5], 150 mM NaCl, 0.5 mM EDTA, and 1% NP-40). Lysates were then mixed with Laemmli buffer and boiled at 95°C for 10 minutes before being centrifuged at 6000 rpm for 5 minutes. Samples were ran on a 4%-12% SDS-PAGE gel and transferred to nitrocellulose membrane (Amersham Biosciences). Immunoblotting was performed using a Western Breeze Chemiluminescence anti-rabbit kit (Invitrogen) according to manufacturer’s instructions. Primary antibody used was anti-GFP polyclonal antibody (Invitrogen) (1:1250).

### Ultrastructure expansion microscopy (UExM)

Ookinetes were allowed to develop in ookinete media for 24 hours before being purified with 13.1-coated beads labelling the protein P28. Once purified, ookinetes were fixed using 4% paraformaldehyde. Fixed cells were attached to 12 mm poly-D-lysine coverslips (A3890401, Gibco) for 30 minutes. Coverslips were then incubated at 4°C overnight in 1.4% formaldehyde and 2% acrylamide in PBS. The following day, gels were prepared using monomer solution (23% sodium acrylate, 10% acrylamide, 0.1% bis-acrylamide in PBS) and setting the monomer solution using 10% APS and 10% TEMED. Gels were incubated for 30 minutes at 37°C. Gels were then denatured for 15 minutes at 37°C and then transferred to Eppendorf tubes and boiled at 95°C for 45 minutes. Gels were then left in distilled water overnight to expand. The next day, gels were washed in PBS for 15 minutes before being incubated in primary antibodies diluted in 3% BSA at 4°C overnight. Gels were then washed three times in 0.1% PBS-Tween, 10 minutes per wash. Gels were then incubated in secondary antibodies for three hours at room temperature before being washed in 0.1% PBST as described. Gels were again left to expand in distilled water overnight. Method adapted from [38].

Primary antibodies used were α-tubulin (Sigma T9026, 1:1000), GFP (Thermo Fisher, 1:250). Secondary antibodies used were mouse Alexa Fluor 488 (1:1000), rabbit Alexa Fluor 568 (1:1000). Stains used were Atto 594 NHS Ester (Merck 08741, 1:200). Gels were imaged in glass bottom dishes (MatTek, P35GC-1.5-14-C). A Zeiss Elyra PS.1-LSM780 microscope was used to image gels. Image analysis was performed in Zeiss Zen 2012 Black edition or ImageJ.

### Structured Illumination Microscopy (SIM)

A small volume of purified schizonts were placed onto a slide and covered with a large coverslip (50 x 34 mm) to create an immobilised monolayer for imaging. SIM imaging was performed on a Zeiss Elyra PS.1-LSM780 microscope fitted with a Plan Apochromat 63x/1.4 oil objective. The correction collar of the objective was set to 0.17 for optimal contrast. Settings used were lasers, 405 nm: 20%, 488 nm: 16%, exposure times, 405 nm: 200 ms, 488 nm: 100 ms, three grid rotations, five phases. Bandpass filters BP 420-480 + LP 750 and BP 495-550 + LP 750 were used for the blue and green channels, respectively.

### Electron microscopy

Ookinetes were purified using 13.1-coated beads and then fixed in 4% glutaraldehyde in 0.1M phosphate buffer and processed for electron microscopy. Briefly, samples were post-fixed in 1% osmium tetroxide in 1.5% potassium ferrocyanide in 0.1 M phosphate buffer for 2 hours at room temperature in the dark. Samples were then incubated in 2% uranyl acetate in ddH_2_O for 90 mins, dehydrated in ethanol (a progressive series of ethanol concentrations from 20%, 40%, 90% to 100% with two changes in absolute ethanol) and embedded in TAAB 812 Hard resin (TAAB, catalogue number T030). Ultrathin sections (approximately 70 nm) were cut from the sample using a diamond knife and mounted on copper grids. The grids were post stained with 5% uranyl acetate followed by lead citrate to enhance contrast. Sections were examined using a Jeol JEM 1400 Flash transmission electron microscope (JEOL) at 120 kV.

### RNA isolation and quantitative real-time PCR (qRT-PCR)

RNA was extracted from purified schizonts and ookinetes using an RNA extraction kit (Stratagene) according to manufacturer instructions. Complementary DNA was made using a High-Capacity RNA-to-cDNA kit (Applied Biosystems) according to manufacturer instructions. Gene expression was quantified from 80 ng total cDNA using SYBR Green Fast Master Mix kit (Applied Biosystems). Primers were designed using Primer3 (Primer-BLAST, NCBI). A 7500 Fast machine (Applied Biosystems) was used to perform the analysis with the following cycling conditions: 95°C for 20 seconds, followed by 40 cycles of 95°C for 3 seconds and 60°C for 30 seconds. Two technical replicates and two biological replicates were performed for each gene tested. Two control reference genes were used: *arginyl-tRNA synthetase* (PBANKA_1434200) and *hsp70* (PBANKA_0818900). Primers used are listed in Supp. Table 3.

### Ookinete motility assay and quantification

Our ookinete motility assay was previously described in Zeeshan *et al*., 2022. Briefly, blood was taken from mice and incubated in double the volume of ookinete media for 24 hours. Cells were resuspended and mixed with Matrigel (Corning) in a 1:1 ratio.

Cells were then added to a microscope slide, a coverslip placed over them and nail polish was used to seal the coverslip to the slide. Slides were allowed to set for 30 minutes before imaging. Time series images were captured on the Zeiss AxioImagermicroscope previously described, using the 63 X objective and DIC imaging. Analysis was performed in ImageJ using the Manual Tracking plug-in. Briefly, a spot was placed at the apical end of the ookinete in all images in a time series. Manual tracking allowed the path taken by the spot to be plotted and measured giving measurements for distance and speed. Distances between frames were added together to give total distance travelled. Measurements for speed were averaged to give mean speed travelled.

### Phosphoproteomics

#### Sample preparation (SDS buffer-FASP procedure)

Cell lysis was performed in 400 μl of 2% SDS, 25 mM NaCl, 50 mM Tris (pH 7.4), 2.5 mM EDTA and 20 mM TCEP supplemented with 1x Halt™ protease and phosphatase inhibitor. Samples were vortexed and then heated at 95°C for 10 min with 400 rpm mixing on a thermomixer. DNA was sheared *via* four sonication pulses of 10 s each at 50% power. Samples were then centrifuged for 30 min at 17’000 g and the supernatant was collected. After protein assay, a SDS-PAGE was performed to assess the quality of the samples and the volume corresponding to 600 μg of protein was taken for each sample, completed to 300 μl with Tris-HCl 0.1M pH 8.5 and of the samples were incubated with 48 μl of iodoacetamide 0.5 M for 1h at room temperature. Protein were digested based on the FASP method [44] using Amicon® Ultra-4, 30 KDa as centrifugal filter units (Millipore). Trypsin (Promega) was added at 1:80 enzyme/protein ratio, and digestion was performed overnight at room temperature. The resulting peptide samples were desalted using a Pierce^TM^ Peptide Desalting Spin Columns (Thermo Fisher Scientific) according to manufacturer’s instruction and then completely dried under speed-vacuum.

#### TMT11plex-labelling procedure

Peptide concentration was determined using a colorimetric peptide assay (Thermo Fisher Scientific). Briefly, 140 μg of each sample was labelled with 800 μg of the corresponding TMT-11plex reagent previously dissolved in 220 μl of 36% CH3CN, 200 mM EPPS (pH 8.5). The reaction was performed for 1 hour at room temperature and then quenched by adding hydroxylamine to a final concentration of 0.3% (v/v). Labelled samples were pooled together, dried under speed-vacuum and desalted using a Pierce^TM^ Peptide Desalting Spin Columns (Thermo Fisher Scientific) according to manufacturer’s instruction and then completely dried under speed-vacuum. The tagging efficiency was around 90%.

#### Phosphopeptide enrichment

Phosphopeptides were enriched using the High-Select Fe-NTA Phosphopeptide Enrichment Kit (Thermo Fisher Scientific) following manufacturer’s instructions. Phosphopeptides fraction as well as flow-through fraction were desalted with Pierce^TM^ Peptide Desalting Spin Columns (Thermo Fisher Scientific) according to manufacturer’s instruction and then completely dried under speed-vacuum.

#### ESI-LC-MSMS

Samples were reconstituted in loading buffer (5% CH3CN, 0.1% FA) and the peptide concentration was determined using a colorimetric peptide assay (Thermo Fisher Scientific). For the flow through (FT) and for the phosphopeptide-enriched fractions, 3 μg were injected on column. LC-ESI-MS/MS was performed on an Orbitrap Fusion Lumos Tribrid mass spectrometer (Thermo Fisher Scientific) equipped with a Vanquish NEO liquid chromatography system (Thermo Fisher Scientific). Peptides were trapped on an Acclaim pepmap100, C18, 3μm, 75μm x 20mm nano trap-column (Thermo Fisher Scientific) and separated on a 75 μm x 500 mm, C18 ReproSil-Pur (Dr. Maisch GmBH), 1.9 μm, 100 Å, home-made column. The analytical separation was run for 180 min using a gradient of H2O/FA 99.9%/0.1% (solvent A) and CH3CN/H2O/FA 80.0%/19.9%/0.1% (solvent B). The gradient was run from 5 % B to 28 % B in 160 min, then to 40% B in 20 min, then to 99% B in 10 min with a final stay of 20 min at 99 % B. Flow rate was of 250 nL/min an total run time was of 210 min. Data-Dependant Acquisition (DDA) was performed with MS1 full scan at a resolution of 120’000 FWHM followed by as many subsequent MS2 scans on selected precursors as possible within 3 second maximum cycle time. MS1 was performed in the Orbitrap with an AGC target of 4x105, a maximum injection time of 50 ms and a scan range from 375 to 1500 *m/z*. MS2 was performed in the Orbitrap at a resolution of 50’000 FWHM using higher-energy collisional dissociation HCD at 38% NCE. Isolation windows was at 0.7 u with an AGC target of 5x104 and a maximum injection time of 86 ms. A dynamic exclusion of parent ions of 60 s. with 10 ppm mass tolerance was applied.

#### Database search

Raw data were processed using Proteome Discoverer 2.4 software (Thermo Fisher Scientific). Briefly, spectra were extracted and searched against the *Plasmodium berghei* ANKA database (PlasmoDB.org, release 68, 4958 entries), the *mus musculus* reference proteome database (UniProt, reviewed, release 2025_08, 17’234 entries) and an in-house database of common contaminant using Mascot (Matrix Science, London, UK; version 2.6.2). Trypsin was selected as the enzyme, with one potential missed cleavage. Precursor ion tolerance was set to 20 ppm and fragment ion tolerance to 0.02 Da. Carbamidomethylation of cysteine (+57.021), TMT6plex (+229.163) on lysine residues and TMT6plex (+229.163) on peptide N-termini were specified as fixed modification. Oxidation of methionine (+15.995) was set as variable modifications, as well as phosphorylation of serine, threonine and tyrosine. The search results were validated with Percolator. PSM and peptides were filtered with a false discovery rate (FDR) of 1%, and then grouped to proteins with again an FDR of 1% and using only peptides with high confidence level. Both unique and razor peptides were used for quantitation and protein and peptides abundances values were based on S/N values of reporter ions. The abundances were normalised on “Total Peptide Amount” excluding phosphopeptides and then scaled with the “On all Average” parameter. The protein ratios were directly calculated from the grouped protein abundances and associated p-values were calculated with ANOVA test based on the Abundances of individual proteins or peptides. The ptmRS mode was used for phosphorylation modifications sites localization and validation. Only phosphosites with a probability of more than 75% are considered as validated.

### Statistical analysis

All graphs and statistical analysis were produced in GraphPad Prism 10. Students t tests were used to assess differences between control and experimental groups. Statistical significance is denoted as: *p<0.05, **p<0.01, ***p<0.001, ****p<0.0001 or ns = non-significant. For qRT-PCR experiments, one-way ANOVA was performed on -ΔCt values from two biological replicates and two technical replicates.

## Author contributions

PS and RT conceived the project. SP, MZ, AM, DB, DSG, AAH, RT performed and analysed live cell, knockdown, proteomics, phosphoproteomics, mosquito and mice work. MH, DSF and SV performed electron microscopy. CH, GK, DV performed ookinete injection experiments, RT, PS, DV, GK, SV coordinated the project. SP, RT and DSG wrote the manuscript and all other reviewed and edited it.

## Acknowledgements

We acknowledge technical assistance from Igor Blatov, Lizzie Ison and Robert Markus at The University of Nottingham and thank Dr Ryuji Yanase for his modified protocol to purify ookinetes for phosphoproteomics and TEM work. We also thank Andrew Bottrill at the University of Warwick for providing proteomic work assistance and Temesgen M Kebede at Imperial College for mouse work. For Open Access, the authors have applied a CC BY public copyright licence to any Author Accepted Manuscript version arising from this submission.

## Supplementary materials

**Supplementary Figure 1 – Generation and characterisation of PK2-GFP construct and transgenic line. (A)** Schematic demonstrating single homologous recombination strategy to make PK2-GFP using the *Pbpk2* locus and a GFP tagging construct. Arrows represent the position of the primers used to confirm integration. **(B)** Diagnostic PCR to confirm correct integration of *pk2* into the GFP construct. **(C)** Western blot to check the molecular weight of the GFP-tagged PK2 protein using an antibody against GFP. WT GFP was used as a control (molecular weight ∼25 kDa). This Western blot has been spliced to remove other samples that were ran on the same gel. **(D)** Live cell imaging of PK2-GFP x MyoAch ookinete crosses. White box indicates zoom of apical tip. Scale bar, 2 µm. **(E)** 3D-SIM imaging of PK2-GFP ookinetes. Scale bar, 2 µm. **(F)** 3D-SIM imaging of PK2-GFP x MyoAch ookinete crosses. Scale bar, 2 µm. **(G)** U-ExM of PK2-GFP merozoites stained with NHS Ester (grey), GFP (green) and Hoechst (blue). Scale bar, 2 µm.

**Supplementary Figure 2 – Generation and characterisation of conditional knockdown PK2 PTD. (A)** Schematic demonstrating double homologous recombination strategy used to swap the promoter of *pk2* for the promoter of *ama1*. Arrows 1 and 2 indicate position of primers used to confirm 5’ integration and 3 and 4 the position of primers used to confirm 3’ integration. **(B)** Diagnostic PCR was performed to confirm correct integration using primers IntPTD 61 5’ and 5’IntPTD to check the integration of the selectable marker and IntPTD61 3’ and 3’ IntAMA1 to check the integration of the *ama1* promoter. **(C)** qRT-PCR analysis of *pk2* transcript levels in WT GFP and *pk2ptd* gametocytes activated for 15 mins. Error bars = ±SEM of two independent experiments **(D)** qRT-PCR analysis of *pk2* transcript levels in WT GFP and *pk2ptd* ookinetes 24 hpa. Error bars = ±SEM of two independent experiments. **(E)** qRT-PCR analyses of the transcript levels of highly expressed invasion genes – CTRP, SOAP and CHT1 – in WT GFP and *pk2ptd* ookinetes 24 hpa. Error bars = ±SEM of two independent experiments. **(F)** qRT-PCR analyses of the transcript levels of less-expressed invasion genes – Celtos and P28 – in WT GFP and *pk2ptd* ookinetes 24 hpa. Error bars = ±SEM of two independent experiments.

**Supplementary Table 1: Proteomics of *pk2ptd* v WT GFP ookinetes 24 hpa**

**Supplementary Table 2: Phosphoproteomics of *pk2ptd* v WTGFP ookinetes 24 hpa**

**Supplementary Table 3 - Primers used in the study**

